# VISTA: Visualizing the Spatial Transcriptome of the *C. elegans* Nervous System

**DOI:** 10.1101/2023.04.28.538711

**Authors:** David Liska, Zachery Wolfe, Adam Norris

## Abstract

**Summary:** Profiling the transcriptomes of single cells without sacrificing spatial information is a major goal of the field of spatial transcriptomics, but current technologies require tradeoffs between single-cell resolution and whole-transcriptome coverage. In one animal species, the nematode worm *C. elegans*, a comprehensive spatial transcriptome with single-cell resolution is attainable using existing datasets, thanks to the worm’s invariant cell lineage and a series of recently-generated single cell transcriptomes. Here we present VISTA, which leverages these datasets to provide a visualization of the worm spatial transcriptome, focusing specifically on the nervous system. VISTA allows users to input a query gene and visualize its expression across all neurons in the form of a “spatial heatmap” in which the color of a cell reports the expression level. Underlying gene expression values (in Transcripts Per Million) are displayed when an individual cell is selected. We provide examples of the utility of VISTA for identifying striking new gene expression patterns in specific neurons, and for resolving cellular identities of ambiguous expression patterns generated from *in vivo* reporter genes. The ability to easily obtain gene-level snapshots of the neuronal spatial transcriptome should facilitate studies on neuron-specific gene expression and regulation, and provide a template for the high-resolution spatial transcriptomes the field hopes to obtain for various animal species in the future.

**Availability and Implementation:** VISTA is freely available at the following URL: https://public.tableau.com/app/profile/smu.oit.data.insights/viz/VISTA_16814210566130/VISTA

## 1 Introduction

The prospect of profiling the entire transcriptome of individual cells without sacrificing positional information is a tantalizing possibility held forth by the emerging field of spatial transcriptomics^1,2^. Although the field is growing rapidly, current platforms entail tradeoffs in which the user must sacrifice either single-cell resolution or whole-transcriptome coverage^3–5^.

The nematode worm *C. elegans* has a history of trailblazing in the field of metazoan single-cell gene expression, providing both spatial and developmental *in vivo* context^6–10^. Because of the invariant cell lineage and stereotyped cell positioning from one individual to the next, a universal spatial “worm atlas” can be constructed with single cell resolution. Once single-cell transcriptomic data is overlaid upon this atlas, the result will be a reconstructed spatial transcriptome that can in principle be applied to each single cell of an entire organism.

Many systematic single-cell transcriptomic datasets have been generated in *C. elegans*, including pan-cellular single-cell RNA Seq^6^, sequencing of individual subsets of cell types throughout embryonic and larval development^7^, and pan-cellular sequencing across embryonic development^11^ and throughout the aging process^12^. Recent work focused on the nervous system has provided single-cell transcriptomes for all neuronal cell types^13^ as well as a complementary “deep transcriptome” for select neuronal cell types obtained by cell-specific FACS sorting^14^.

To integrate existing neuronal single-cell transcriptomes with spatial cellular information, we present VISTA-a user-friendly web browser application allowing users to visually interrogate gene expression across the entire *C. elegans* nervous system. VISTA accepts gene queries in a variety of formats and returns single-cell gene expression heatmaps. The first is an alphabetically-ordered heatmap in which individual tiles represent neuronal cell types. The second is a spatial heatmap in which individual neurons are displayed in their *in vivo* anatomical positions and colored according to their scaled gene expression levels.

## 2 Implementation

VISTA visualizations were created with Tableau and are freely available on Tableau Public. The anatomical imagery was constructed as a profile snapshot of OpenWorm’s (openworm.org) 3D model. Spatial cell position information was derived from WormAtlas (wormatlas.org) and OpenWorm through landmark identification and relative placement. Single-cell gene expression data was obtained from two datasets generated by the CeNGEN consortium^13–15^. Single-cell transcripts per million (TPM) values were obtained directly from published data^13^. For the bulk sorted data, we first normalized using DESeq^16^ (default normalization parameters/estimateSizeFactors) then calculated TPM for each cell type. The Single-cell TPM values and bulk sorted data were combined by a union before being joined on the neuron identifier to manually generated spatial data set and curated descriptive neuron information.

## 3 Usage

To initialize a VISTA spatial search, a gene ID is input into the search bar. Three gene ID formats are supported: common names, sequence names, and Wormbase gene IDs. The output is two forms of heatmap (Figure 1A). In the spatial heatmap, individual neurons are displayed in their *in vivo* anatomical positions, and colored according to their scaled gene expression levels. Hovering the cursor over an individual neuron reveals the TPM value for the queried gene in that neuron. In the alphabetical heatmap, individual tiles represent neuronal cell types, and are ordered alphabetically. Hovering the cursor over the tile reveals the numerical TPM value, and clicking the tile highlights the location of the neuron on the spatial heatmap.

**Figure 1:**
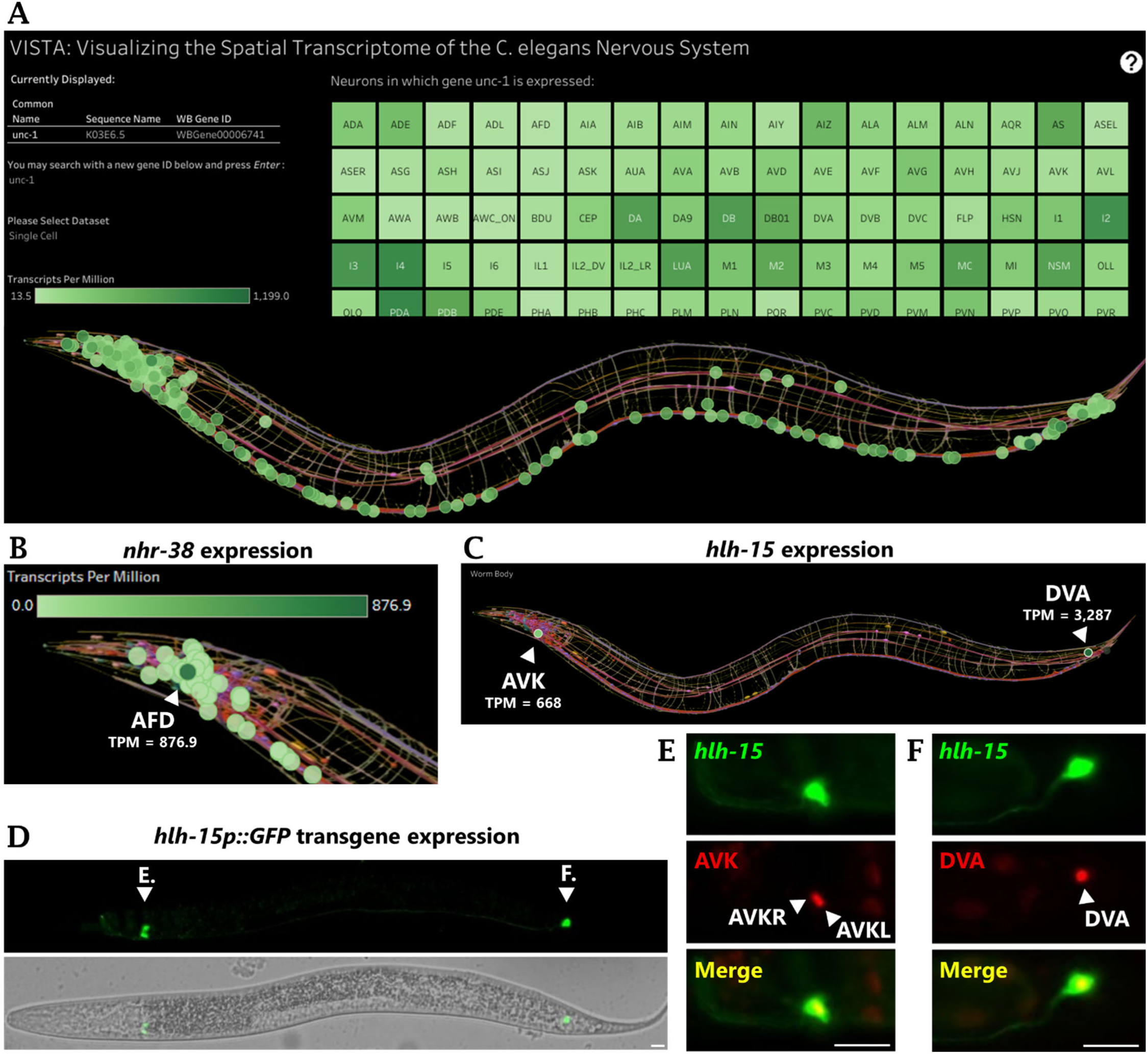
VISTA visualizes spatial transcriptomic gene expression data with single-cell resolution. (A) The VISTA display of spatial gene expression data for the gene *unc-1*. Alphabetical heatmap with individual neuron tiles are above, and spatial heatmap with each cell placed in its anatomical position is below. Darker colors represent cells with higher expression. (B) Highly specific expression of *nhr-38* in a single neuron, the AFD neuron (bulk-sorted data displayed). (C) *hlh-15* is predicted to be highly expressed in a single paired head neuron (AVK) and a single tail neuron (DVA). Single-cell data displayed. (D) *hlh-15p::GFP* transgene reveals expression in a single paired head neuron and a single tail neuron. Letters E. and F. indicate the neurons imaged at higher resolution in the respective figure panels (not from the same worm.) (E) *hlh-15p::GFP*-expressing head neurons perfectly overlap with AVK nuclear mCherry driven by *flp-1* promoter. (F) *hlh-15p::GFP*-expressing tail neuron perfectly overlap with DVA nuclear mCherry driven by *nlp-12* promoter. Scale bars represent 10 μm.

Users can zoom, pan, and select cells of interest using an instrument panel located at the lower left-hand corner of the dashboard. Users can toggle back and forth between single-cell and bulk sorted datasets using the drop-down menu. The two datasets are complementary in a number of ways: the single-cell dataset covers a much larger number of neuron types, while the bulk sorted dataset is able to distinguish between two classes of neurons that are indistinguishable in the single cell data (VD versus DD motor neurons). The bulk dataset is better suited to detect lowly-expressed genes and mRNAs that are not polyadenylated, but is also subject to low levels of contaminating cells and thus gene expression signatures^14^. As such it can be useful to query both datasets when possible.

## 4 Use Case

As an example of the utility of VISTA, we investigated whether we could predict interesting new spatially-restricted gene expression patterns simply by visual browsing through genes on VISTA. We focused on transcription factor genes, which have been extensively studied for their cell-specific expression and function and thus represent good “ground truth” candidates^17–20^. We validated many such ground truth TFs, for example expression of nuclear hormone receptor TFs to single neurons: *nhr-38* exclusive to the AFD thermosensory neuron^21^ (Figure 1B), and *odr-7* exclusive to the AWA chemosensory neuron^22^.

We also identified a striking expression pattern for the helix-loop-helix TF *hlh-15*, in which strong expression is confined to a single neuron in the tail (DVA) and a single paired neuron in the head (AVKL/R) (Figure 1C). A previous study generated a promoter::GFP transgene for *hlh-15* and likewise reported expression in a single tail neuron and a single paired head neuron,^23^ but tentatively assigned the expression to the DVA and RIF neurons based on cell position and morphology. To clarify the discrepancy, we obtained the *hlh-15p*::GFP transgene (Figure 1D) and crossed it with nuclear red fluorescent transgenes (H2B::mCherry) expressed in only the DVA neuron or only the AVK neuron^24^. We found that the DVA mCherry marks the nucleus of the cell identified in the tail, and the AVK mCherry marks the nucleus of the paired head neurons (Fig 1E-F), confirming VISTA’s cell-type identification for *hlh-15* gene expression.

This use case highlights the value of VISTA spatial transcriptomic visualization in identifying new interesting expression patterns and in clarifying known expression patterns. In the case of *hlh-15*, the spatial transcriptome reveals a notable symmetry, as it is expressed in a single head neuron that sends a neurite posteriorly all the way to the tail, and a single tail neuron that sends a neurite all the way anterior to the head.

## 5 Conclusion

VISTA enables visualization of the spatial transcriptome with single-cell resolution across the entire nervous system of an animal for the first time. This spatial atlas in *C. elegans* provides a template for the sort of resolution the field of spatial transcriptomics hopes to attain for various animal species in the future. The VISTA platform should be amenable to additional experimental variations such as spatially-resolved gene expression changes in mutant animals^25^ or animals exposed to specific environmental or experimental conditions^26^. Likewise, additional tissues and cell types can be incorporated using existing pan-cellular datasets^6^. Finally, VISTA should be amenable to the incorporation of additional types of gene expression data, for example the spatial visualization of alternative splicing events in single cells^27,28^ should be displayable using the same format and datasets described here.

## Funding

National Institute of General Medical Sciences of the National Institutes of Health [R35GM133461]

## Data Availability

VISTA is available at the URL https://public.tableau.com/app/profile/smu.oit.data.insights/viz/VISTA_16814210566130/VISTA. The hosted Tableau workbook is freely downloadable and can be used to locally share and interact with the data using the free Tableau Reader software https://www.tableau.com/products/reader. Underlying gene expression data is included here (Supplementary Table 1-2), is available on GitHub https://github.com/xcwolfe/VISTA-C-elegans and the original raw data is available in the original single-cell transcriptome publications^13,14^.

## Notes

https://public.tableau.com/app/profile/smu.oit.data.insights/viz/VISTA_16814210566130/VISTA

### Competing Interest Statement

The authors have declared no competing interest.

### Summary of Updates

Fixed author affiliation mis-spell

## References

1. Williams, C. G., Lee, H. J., Asatsuma, T., Vento-Tormo, R. & Haque, A. An introduction to spatial transcriptomics for biomedical research. Genome Med 14, 68 (2022).

2. Marx, V. Method of the Year: spatially resolved transcriptomics. Nat Methods 18, 9–14 (2021).

3. Eng, C.-H. L. et al. Transcriptome-scale super-resolved imaging in tissues by RNA seqFISH. Nature 568, 235–239 (2019).

4. Larsson, L., Frisén, J. & Lundeberg, J. Spatially resolved transcriptomics adds a new dimension to genomics. Nat Methods 18, 15–18 (2021).

5. Ståhl, P. L. et al. Visualization and analysis of gene expression in tissue sections by spatial transcriptomics. Science 353, 78–82 (2016).

6. Cao, J. et al. Comprehensive single-cell transcriptional profiling of a multicellular organism. Science 357, 661–667 (2017).

7. Spencer, W. C. et al. A spatial and temporal map of C. elegans gene expression. Genome Res 21, 325–341 (2011).

8. Dupuy, D. et al. Genome-scale analysis of in vivo spatiotemporal promoter activity in Caenorhabditis elegans. Nat Biotechnol 25, 663–668 (2007).

9. Sulston, J. E. & Horvitz, H. R. Post-embryonic cell lineages of the nematode, Caenorhabditis elegans. Dev Biol 56, 110–156 (1977).

10. Chalfie, M., Horvitz, H. R. & Sulston, J. E. Mutations that lead to reiterations in the cell lineages of C. elegans. Cell 24, 59–69 (1981).

11. Packer, J. S. et al. A lineage-resolved molecular atlas of C. elegans embryogenesis at single-cell resolution. Science 365, eaax1971 (2019).

12. Roux, A. E. et al. The complete cell atlas of an aging multicellular organism. http://biorxiv.org/lookup/doi/10.1101/2022.06.15.496201 (2022) doi:10.1101/2022.06.15.496201.

13. Taylor, S. R. et al. Molecular topography of an entire nervous system. http://biorxiv.org/lookup/doi/10.1101/2020.12.15.422897 (2020) doi:10.1101/2020.12.15.422897.

14. Barrett, A. et al. Integrating bulk and single cell RNA-seq refines transcriptomic profiles of specific C. elegans neurons. http://biorxiv.org/lookup/doi/10.1101/2022.04.05.487209 (2022) doi:10.1101/2022.04.05.487209.

15. Hammarlund, M., Hobert, O., Miller, D. M. & Sestan, N. The CeNGEN Project: The Complete Gene Expression Map of an Entire Nervous System. Neuron 99, 430–433 (2018).

16. Love, M. I., Huber, W. & Anders, S. Moderated estimation of fold change and dispersion for RNA-seq data with DESeq2. Genome Biology 15, (2014).

17. Hobert, O., Glenwinkel, L. & White, J. Revisiting Neuronal Cell Type Classification in Caenorhabditis elegans. Curr. Biol. 26, R1197–R1203 (2016).

18. Reilly, M. B., Cros, C., Varol, E., Yemini, E. & Hobert, O. Unique homeobox codes delineate all the neuron classes of C. elegans. Nature 584, 595–601 (2020).

19. Thompson, M. et al. Splicing in a single neuron is coordinately controlled by RNA binding proteins and transcription factors. eLife 8, e46726 (2019).

20. Taylor, M., Marx, O. & Norris, A. D. TDP-1 and FUST-1 co-inhibit exon inclusion and control fertility together with transcriptional regulation. http://biorxiv.org/lookup/doi/10.1101/2023.04.18.537345 (2023) doi:10.1101/2023.04.18.537345.

21. Miyabayashi, T., Palfreyman, M. T., Sluder, A. E., Slack, F. & Sengupta, P. Expression and function of members of a divergent nuclear receptor family in Caenorhabditis elegans. Dev Biol 215, 314–331 (1999).

22. Sengupta, P., Colbert, H. A. & Bargmann, C. I. The C. elegans gene odr-7 encodes an olfactory-specific member of the nuclear receptor superfamily. Cell 79, 971–980 (1994).

23. Grove, C. A. et al. A multiparameter network reveals extensive divergence between C. elegans bHLH transcription factors. Cell 138, 314–327 (2009).

24. Wang, H. et al. cGAL, a temperature-robust GAL4-UAS system for Caenorhabditis elegans. Nat Methods 14, 145–148 (2017).

25. Liang, X., Calovich-Benne, C. & Norris, A. Sensory neuron transcriptomes reveal complex neuron-specific function and regulation of mec-2/Stomatin splicing. Nucleic Acids Res 50, 2401–2416 (2022).

26. Harris, N. et al. Molecular encoding of stimulus features in a single sensory neuron type enables neuronal and behavioral plasticity. Curr Biol S0960-9822(23)00263–4 (2023) doi:10.1016/j.cub.2023.02.073.

27. Liang, X., Taylor, M., Napier-Jameson, R., Calovich-Benne, C. & Norris, A. A Conserved Role for Stomatin Domain Genes in Olfactory Behavior. eNeuro 10, ENEURO.0457-22.2023 (2023).

28. Norris, A. D. et al. A Pair of RNA-Binding Proteins Controls Networks of Splicing Events Contributing to Specialization of Neural Cell Types. Molecular Cell 54, 946–959 (2014).

